# Collective nuclear behavior shapes bilateral nuclear symmetry for subsequent left-right asymmetric morphogenesis in *Drosophila*

**DOI:** 10.1101/2020.10.15.340521

**Authors:** Dongsun Shin, Mitsutoshi Nakamura, Yoshitaka Morishita, Mototsugu Eiraku, Tomoko Yamakawa, Takeshi Sasamura, Masakazu Akiyama, Mikiko Inaki, Kenji Matsuno

## Abstract

Proper organ development often requires nuclei to move to a specific position within the cell. To determine how nuclear positioning affects left-right (LR) development in the *Drosophila* anterior midgut (AMG), we developed a surface-modeling method to measure and describe nuclear behavior at stages 13-14, captured in three-dimensional time-lapse movies. We describe the distinctive positioning and a novel *collective nuclear behavior* by which nuclei align LR-symmetrically along the anterior-posterior axis in the visceral muscles that overlie the midgut and are responsible for this organ’s LR-asymmetric development. Wnt4 signaling is crucial for the collective behavior and proper positioning of the nuclei, as are myosin II and LINC complex, without which the nuclei failed to align LR-symmetrically. The LR-symmetric positioning of the nuclei is important for the subsequent LR-asymmetric development of the AMG. We propose that the bilaterally symmetrical positioning of these nuclei may be mechanically coupled with subsequent LR-asymmetric morphogenesis.

## Introduction

Directional left-right (LR) asymmetry, which is evident in many animals’ external and internal morphology, is genetically determined^1–5^. Recent studies show that the mechanisms determining LR-asymmetry are evolutionarily divergent^6–9^. In vertebrates, several different mechanisms contribute to LR-asymmetric development, including nodal flow, LR-asymmetric proton influx, and LR-asymmetric cell migration; some of these mechanisms have parallel functions^9–10^. In Lophotrochozoa and Ecdysozoa, intrinsic cell chirality plays a key role in LR-asymmetric development. For example, cell chirality in snail and nematode blastomeres determines their subsequent LR-asymmetric organ and body development^1,5,11^. In *Drosophila*, the LR-asymmetrical development of several organs also relies on cell chirality, which is controlled by the myosin 1D gene^3,12–16^. Importantly, chiral cells are also found in vertebrates and are thought to contribute to their LR-asymmetric development^17–18^. However, the molecular mechanisms of LR-asymmetric development in invertebrates remain largely unclear. *Drosophila* is an excellent model system for studying these mechanisms^4,7,19^.

At least one other mechanism besides cell chirality is responsible for creating LR-asymmetry in *Drosophila*^20–22^. The first detectable LR-asymmetry in the developing *Drosophila* anterior midgut (AMG), observed in the visceral muscles overlying the epithelial tube of the midgut, occurs independently of cell chirality^20–22^. Initially, the long axis of nuclei in these visceral muscle cells is aligned perpendicular to the midline; however, this angle changes and becomes LR-asymmetrical in ventral-region nuclei at stage 13-14, just before overall LR-asymmetric morphological changes begin^20–22^. These visceral muscles play a crucial role in AMG LR-asymmetric development^20–22^. We previously showed that when the long axis of the nuclei failed to undergo this asymmetric rearrangement (due to augmented JNK signaling or reduced Wnt signaling in the visceral muscles), LR-asymmetry of the AMG also failed^20,22^. We also showed that myosin II (MyoII) is essential for both LR-asymmetric AMG development and the LR-asymmetric rearrangement of the long axis of the nuclei in the visceral muscles, which suggests that the change in the angle of the axis is controlled mechanically^21^. However, the dynamics and underlying mechanisms of this rearrangement remain elusive.

The location of the nucleus, which is the cell’s largest organelle, changes as needed for various cellular contexts and functions^23^. For example, to permit efficient cell migration, the nucleus remains behind the center of the cell, away from the leading edge^24^. The position of the nucleus can differ with tissue morphology and integrity^25–26^, and defects in nuclear positioning are connected with muscular dystrophy and centronuclear myopathy in humans^27–28^. Nuclear migration events depend on LINC (linker of nucleoskeleton and cytoskeleton) complex, which physically links nuclei and F-actin/microtubules^29^.

Here, we studied the movement of nuclei in the visceral muscle overlying the midgut in stage 13-14 wild-type *Drosophila* using three-dimensional (3D) time-lapse movies and quantitative imaging analysis. We found that the nuclei of the visceral muscles were positioned LR-symmetrically in distinct regions along the anterior–posterior axis in wild-type embryos; we refer to this distribution as *proper nuclear positioning* (PNP) hereafter. The densely crowded nuclei in these regions actively rearranged their positions relative to neighboring nuclei; we refer to this as *collective nuclear behavior* (CNB) hereafter. Dally-like protein (Dlp), a component of Wnt signaling, was essential for both PNP and CNB. MyoII and LINC complex were required for PNP but not for CNB. Unexpectedly, however, the nuclei aligned LR-asymmetrically in mutants with disrupted MyoII or LINC complex, although the AMG developed LR-symmetrically. Our results show that the positioning of the nuclei in the visceral muscles is accomplished via multiple regulatory machineries, including Wnt signaling, MyoII, and LINC complex, and that the LR-symmetric positioning of the nuclei is important for the LR-asymmetric development of the AMG.

## Results

### Visceral muscle-cell nuclei are collectively aligned in distinct regions in the wild-type embryonic midgut

The midgut is composed of the epithelial tube and the overlying visceral muscles (Fig. 1a). The visceral muscle cells, which are binucleated and bipolar, align LR-symmetrically at the lateral sides of the embryo with the long axis of each nucleus perpendicular to the midline (Fig. 1a, b)^22,30–31^. In stages 13-14, the leading edges of the visceral muscles extend dorsally and ventrally toward the dorsal and ventral midlines, respectively, and eventually merge at the midlines at late stage 14 (Fig. 1a)^22^. Studies show that the first detectable LR-asymmetry in the AMG is a difference between the right and left sides in the angle between the long axis of the nuclei and the midline in the ventral side of this organ at stage 14^20–22^. Since these studies were conducted in fixed embryos, the events leading to the LR-asymmetry of the visceral muscle nuclei and the AMG are still unclear.

**Fig. 1.**
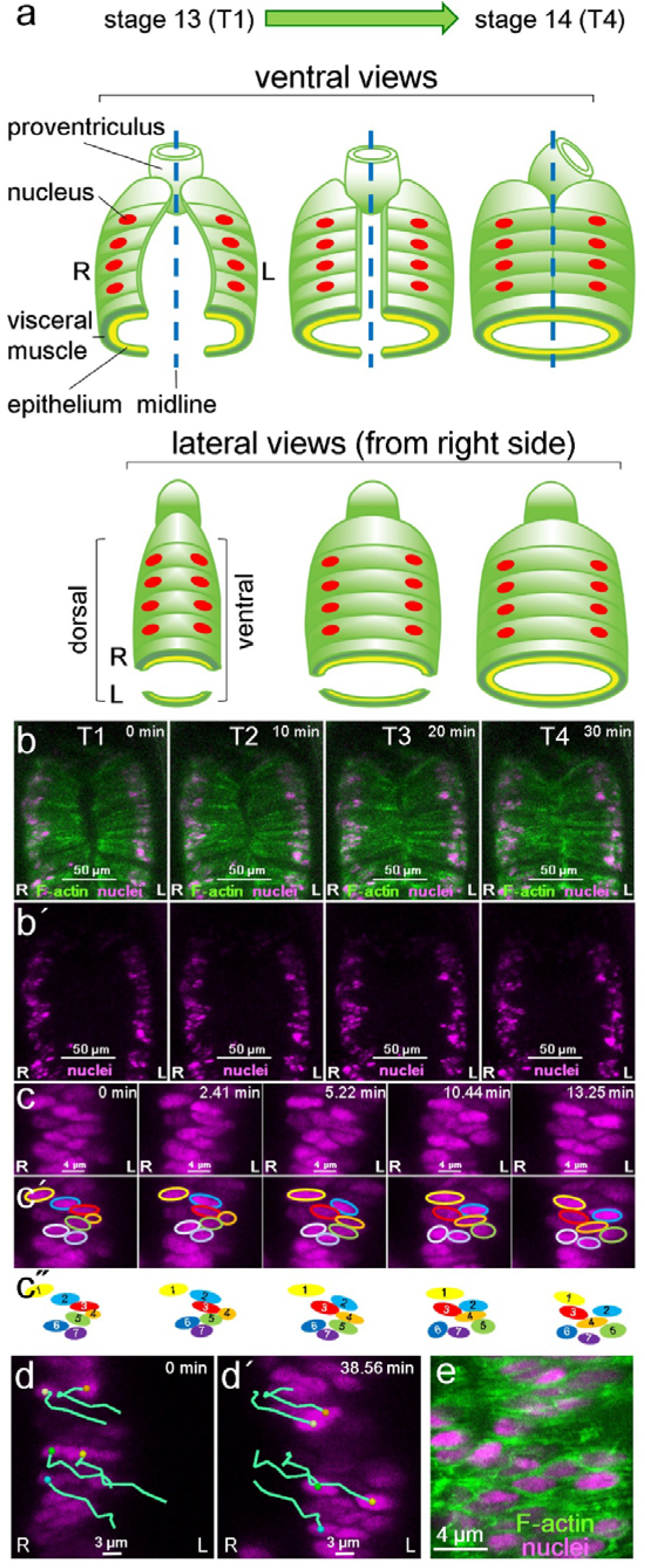
Collective positioning of nuclei in the midgut visceral muscle of wild-type *Drosophila* embryos. **a** Diagram of AMG development from stage 13 (T1) to 14 (T4), showing the epithelium (yellow) and overlying visceral muscles (green) of the midgut in ventral (upper panels) and lateral (lower panels) views. The visceral muscles are binuclear cells that are aligned at the lateral sides at stage 13 (red ovals are nuclei); their leading edges extend toward and eventually merge at the midlines (dotted blue lines) at late stage 14. R, right; L, left. **b, b′** Snapshots from 3D time-lapse movies show ventral views of visceral muscle cells (outlined in green in top panel; F-actin) and nuclei (magenta) at 10-min intervals from T1 (0 min) to T4 (30 min). Scale bars=50 μm. **c** Magnified views of the ventral region, where nuclei (magenta) are densely aligned along the anterior and posterior axes, from snapshots at intervals of 2.41 min beginning at T1. Scale bars=4 μm. **c′** Each nucleus that was visible in all snapshots is outlined in a different color. **c** Colored ovals represent the relative positions of the individual nuclei outlined by the same color in **c′**). **d, d′** Snapshots show right-side visceral muscles. Blue-green lines trace the migration of individual nuclei (magenta) according to their positions at 2.41-min intervals from 0 (T1) to 38.56 min. Colored dots show the position of the center of the nucleus at each time point. Scale bar=3 μm. **e** A magnified view of fixed ventral visceral muscles in which nuclei (magenta) and F-actin (green) were detected by immunostaining. Scale bar=4 μm.

Therefore, to examine the process by which nuclei are arranged, we obtained 3D time-lapse movies of the midgut in developing embryos from stage 13 to 14 using a confocal laser scanning microscope. We used the GAL4/UAS system to drive the visceral muscle–specific expression of *UAS-RedStinger*, which encodes a nuclear DsRed, and of *UAS-lifeact-EGFP*, which encodes a GFP with an actin-binding peptide^32^. The time-lapse movies were obtained from the ventral side of the embryo (Fig. 1b, b′). We designated the time point when the leading edges of the visceral muscles merged at the midline (approximately corresponding to the end of stage 14) as T4; we set T1, T2, and T3 at 30, 20, and 10 min before T4, respectively (Fig. 1b, b′).

In wild-type embryos, the nuclei were densely aligned in distinct regions along both sides of the anterior–posterior axis, creating a region visually similar to the mammalian rib cage, from T1 to T4, in all cases examined (N=10) (Fig. 1b, b′). Collectively, the PNP, referring to the overall positioning of the nuclei with respect to the midline, was maintained from T1 to T4; however, the positions of the individual nuclei changed relative to one another (Fig. 1b, b′). By tracking the position of individual nuclei over time, we found that the nuclei actively moved and adjusted their position relative to each other in all wild-type embryos examined (N=10) (Fig. 1c, c′, c″). We plotted the position of individual nuclei every 2.41 min, starting at T1; at higher magnification, the time-lapse images revealed small movements of the individual nuclei along disparate paths (blue lines in Fig. 1d). Furthermore, despite the dense grouping of the nuclei, they were clearly separated from each other by F-actin along the anterior–posterior axis (Fig. 1e). Thus, the changes in the relative positions of the nuclei were due to the movement of the nuclei within the cells, rather than the rearrangement of entire muscle cells, and we defined this novel collective positioning behavior as CNB. We speculated that, as with other specific nuclear behaviors, CNB is under the control of genetic pathways and may contribute to the LR-asymmetric development of the embryonic midgut^23–28^.

### *dlp* is required in midgut visceral muscles to activate Wnt signaling, which is essential for AMG LR-asymmetry

We conducted a genetic screen that identified a new allele, *dlp^3^*, as a mutation that affects the LR-asymmetric development of the AMG (Fig. 2a, b) (the genetic screen will be reported elsewhere). Our sequence analysis revealed that *dlp^3^* carries a nonsense mutation that introduces a stop codon at the 133^rd^ amino acid residue. Embryos homozygous for *dlp^3^* or *dlp^MH20^* (an amorphic *dlp* allele) or trans-heterozygous for *dlp^3^* and *dlp^MH20^* showed similar defects in AMG LR-asymmetry, including inverted LR-asymmetry and bilateral symmetry (Fig. 2c). The *dlp* gene encodes a core protein of *Drosophila* glypicans, a family of heparan sulfate proteoglycans^33–34^ (including Dlp) that is involved in regulating several cell-signaling pathways, including the Wnt, transforming growth factor-β, and fibroblast growth factor signaling pathways^34–35^.

**Fig. 2.**
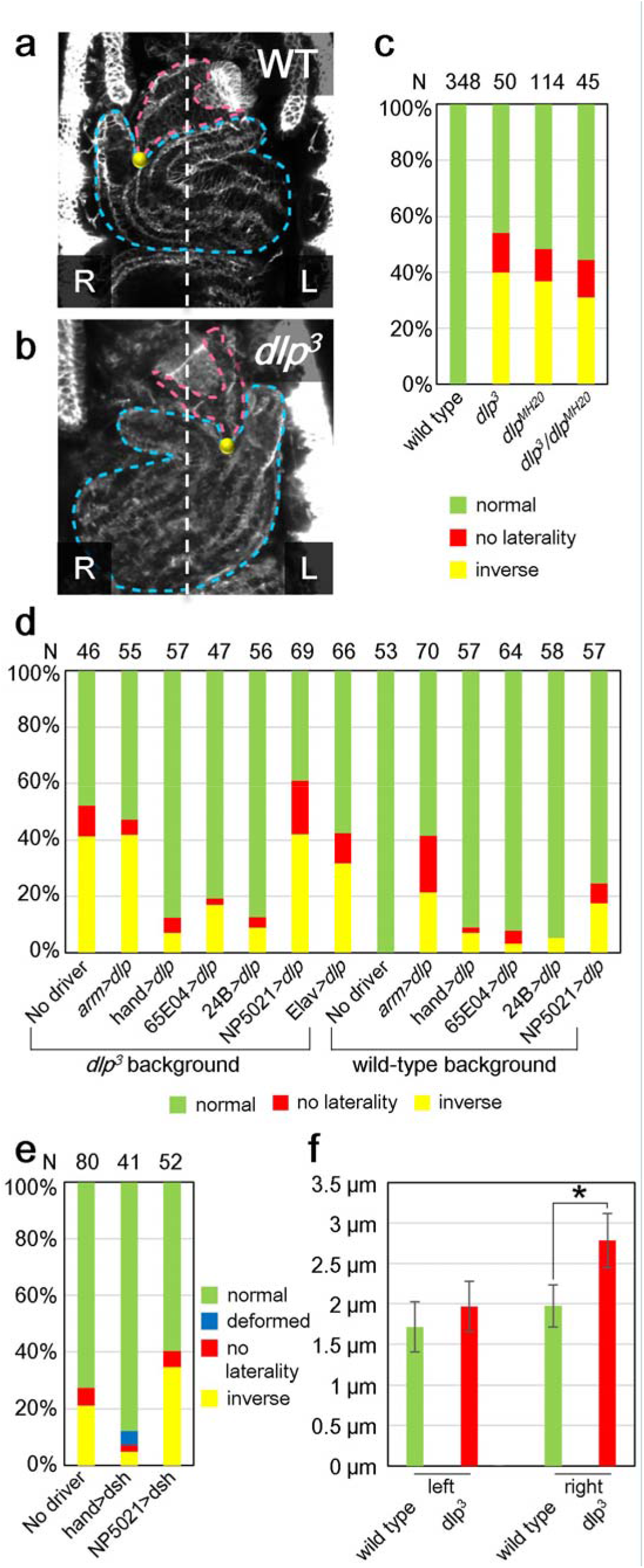
The *dlp* gene is required in the visceral muscles of the midgut to activate Wnt signaling, which is essential for AMG LR-asymmetric development. **a, b** AMG LR-asymmetry in **a)** wild-type (WT) and **b)** *dlp^3^* homozygous embryos viewed from the ventral side, showing the proventriculus (magenta outline), the AMG (blue outline), their connection (yellow spot), and the midline (white outline). L, left, R, right. **c** Bars show the percentage of wild-type, *dlp^3^* homozygous (*dlp^3^*), *dlp^MH20^* homozygous (*dlp^MH20^*), and *dlp^3^/dlp^MH20^* embryos exhibiting an AMG LR phenotype of normal (green), no laterality (red), or inverse (yellow). The number of embryos examined (N) is shown over each bar. **d** Bars show the percentage of embryos from a wild-type or *dlp^3^* homozygous background with an AMG LR phenotype of normal (green), no laterality (red), or inverse (yellow) when *UAS-dlp* was overexpressed by the ubiquitous *Gal4* driver *arm-Gal4* (arm>dlp); by *hand-Gal4* (*hand>dlp*) or *65E04* (*65E04>dlp*), which are specific to circular visceral muscles; *24B* (*24B>dlp*), which is specific to all muscles; *NP5021* (*NP5021>dlp*), which is specific to endodermal epithelium; *Elav-Gal4* (*Elav>dlp*), a pan-neuronal driver; or negative control (No driver). The number of embryos examined (N) is shown over each bar. **e** Bars show the percentage of *dlp^3^* homozygous embryos with an AMG LR phenotype of normal (green), deformed (blue), no laterality (red), or inverse LR (yellow) when *UAS-dsh* was overexpressed with *hand-Gal4* (*hand>dsh*), *NP5021* (*NP5021>dsh*), or negative control (No driver). The number of embryos examined (N) is shown above each bar. **f** Bars indicate the average migration distance (μm) and standard deviation for nuclei in the left and right visceral muscles of wild-type (green) and *dlp^3^* homozygous (red) embryos (N=3 each) over a period of 10 min. *p<0.05.

We previously showed that Wnt4 signaling must be active in the visceral muscle of the AMG for this organ to develop proper LR asymmetry^22^. Thus, we speculated that LR-asymmetric AMG development also requires *dlp* function in the visceral muscle. To test this possibility, we overexpressed *UAS-dlp* specifically in the visceral muscles of the midgut, using the GAL4/UAS system driven by *hand, 65E04*, or *24B*, to see whether it could rescue LR defects in *dlp^3^* homozygotes^22^. Control embryos carrying only *UAS-dlp* (no driver) showed LR defects of the AMG (52% frequency), as did *dlp^3^* homozygotes (54%) (Fig. 2d)^22^. As expected, *UAS-dlp* overexpression markedly suppressed these LR defects when driven by *hand* (frequency of LR defects 12%), *65E04* (19%), or *24B* (12%) (Fig. 2d). In contrast, the frequency of LR defects was not suppressed by overexpressing *UAS-dlp* in the midgut epithelium (*NP5021*, 60%) or nervous system (*Elav-Gal4*, 42%), when compared with control (Fig. 2d). Although *arm-GAL4* is used to drive ubiquitous expression, including in visceral muscles, *dlp* expression driven by *arm-GAL4* in *dlp^3^* homozygotes did not suppress LR defects (Fig. 2d). We speculated that this might be due to potential LR defects associated with *dlp* misexpression in some tissues. Indeed, the ubiquitous misexpression of *UAS-dlp* driven by *arm-Gal4* in wild-type *Drosophila* causes LR defects, whereas control embryos carrying *UAS-dlp* but no *Gal4* driver had no LR defects (Fig. 2d). Taken together, our results show that wild-type *dlp* is required in the visceral muscles for normal LR-asymmetric development of the AMG; which is consistent with our previous finding that normal LR-asymmetric AMG development requires activated Wnt4 signaling in the visceral muscles of the midgut^22^.

Furthermore, recent studies show that Dlp associates with Wnt4 and regulates Wnt signaling in germline cells^36–37^. Therefore, we hypothesized that *dlp* contributes to Wnt4 signaling in the visceral muscles, and thus contributes to LR-asymmetric AMG morphogenesis. To test this possibility, we specifically overexpressed *UAS-disheveled (dsh)*, which can induce Wnt signaling, in the visceral muscles (driven by *hand*) or midgut epithelium (driven by *NP5021*) of *dlp^3^* homozygotes, and examined the effect on LR defects (Fig. 2e)^38–39^. Compared to control embryos (carrying UAS-*dsh* but no *Gal4* driver), the frequency of LR defects associated with the *dlp^3^* mutant decreased when *UAS-dsh* was overexpressed in the visceral muscle (12%) but not when overexpressed in the midgut epithelium (40%) (Fig. 2e). Taken together, these findings suggest that *dlp* is required for the activation of Wnt4 signaling in the visceral muscles, and this activation is essential for normal AMG LR-asymmetric development.

Wnt signaling plays multiple roles in embryonic development^40–41^. Thus, mutants of genes that encode the core components of Wnt signaling show a broad range of phenotypes, including gut deformation, in addition to defects in LR asymmetry^41–42^. Nonetheless, the structure of the midgut in *dlp^3^* mutants was largely normal except for LR randomization, suggesting a specific function for *dlp* in LR-asymmetric morphogenesis (Fig. 2a-c). For example, the extension of the leading edge of the midgut visceral muscles toward the midline is normal in *dlp* mutant embryos, demonstrating that *dlp* is dispensable for this extension (Fig. 3a, b). Therefore, in the following studies of nuclear behavior in AMG visceral muscles, we used the *dlp^3^* mutant to study the visceral muscle–specific depletion of Wnt4 signaling.

**Fig. 3.**
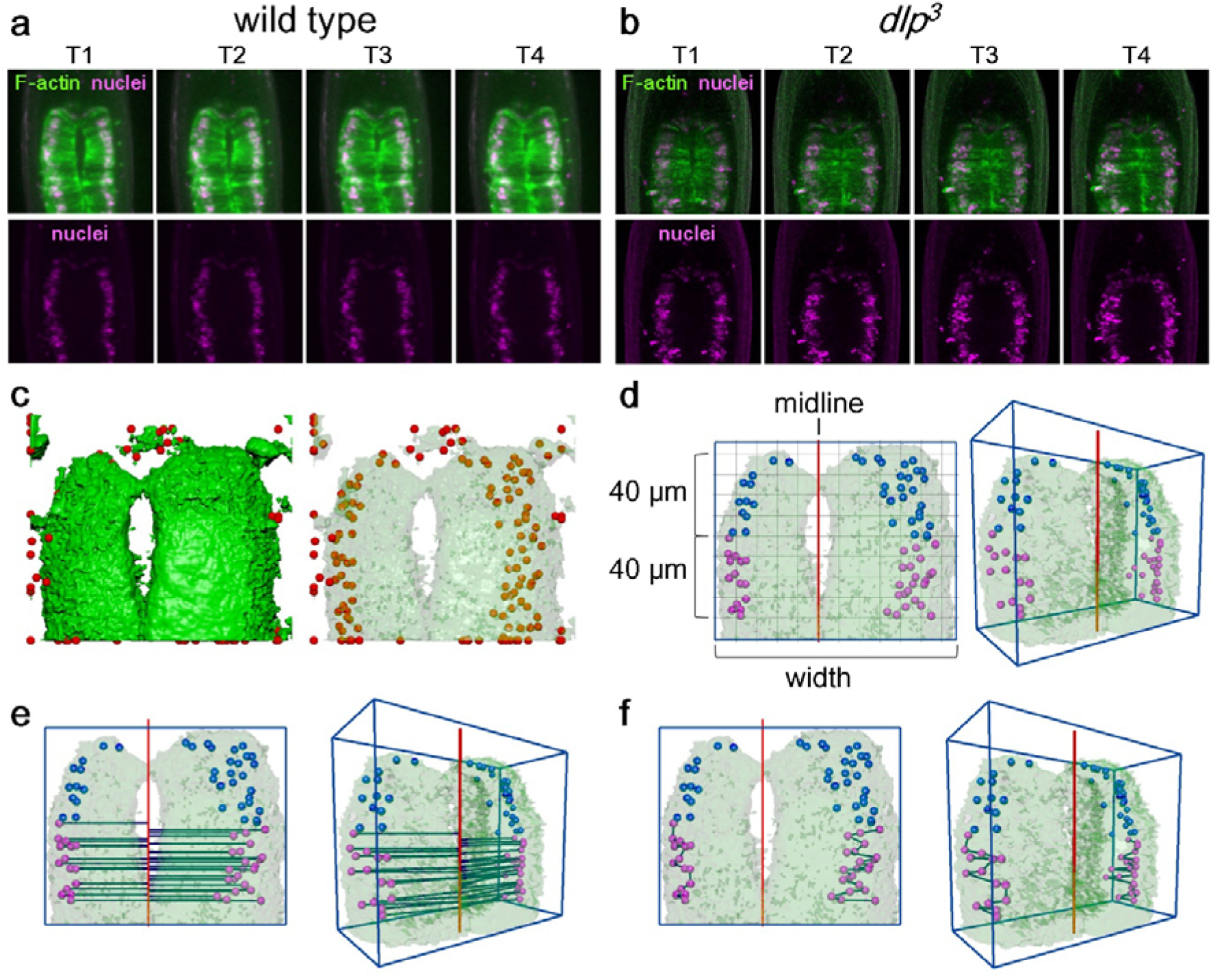
Constructing 3D surface models of the AMG. **a, b** Snapshots from 3D time-lapse movies of the AMG from T1-T4, as viewed from the ventral side in **a**) wild-type (WT) and **b**) *dlp^3^* homozygous embryos. Top panels show nuclei (magenta) and F-actin (green). Lower panels show only nuclei. **c** 3D time-lapse images, as shown in **a** and **b,** were used to reconstruct surface models representing the outer surface of the visceral muscles (green) and the position of nuclei (red) using Imaris software. The surface of the visceral muscles is semitransparent in the model on the right. **d** Assigning the midline in the surface model: surface models of the visceral muscle and nuclei were imported into Maya. A cuboid (blue lines) was computationally configured with its vertical axis set to the length of the embryo’s anterior-posterior axis and its width set to the maximal width of the AMG (shown as width). The midline of the AMG (red line) was determined in 3D space as a line that is parallel to the anterior–posterior grids of the cuboid and connects the merged points of surface-modeled visceral muscles from the left and right sides at T4. Among the nuclei placed in the 3D-surface model, those located 40–80 μm from the anterior tip of the midgut (magenta dots) were selected for further analysis. **e** In the 3D-surface model, a line was drawn from the center of each nucleus to the midline, meeting the midline at right angles (red) in 3D space; the length of the connecting line was automatically measured to obtain the distance between the nucleus and the midline. **f** In the 3D-surface model, a line was automatically drawn in 3D space between each nucleus and its next-most-posterior neighbor. The length of the connecting line was calculated to obtain the distance between the nuclei. Distances between nuclei were measured separately for the right and left sides. **d-f show** dorsal (left) and directional (right) views.

### Wnt4 signaling controls the distance between the nuclei and the midline

To reveal potential defects in the positioning of visceral muscle nuclei in *dlp* mutant embryos, we examined 3D time-lapse movies of the AMG of wild-type and *dlp^3^* homozygous embryos from T1 to T4. When examining 2D snapshots projected from the 3D time-lapse movies, we noticed that the nuclei were more dispersed in *dlp^3^* mutants than in wild-type embryos (Fig. 3a, b). To track the behavior of nuclei in the visceral muscles in the midgut, which is a thick, rounded organ, we took a surface-modeling approach (Fig. 3c, d, e, f). In the surface-modeling analyses, visceral muscles are outlined in green, representing the outer surface of lifeact-EGFP distribution driven by *65E04-Gal4*, a visceral muscle–specific *Gal4* driver (Fig. 3c, left). Nuclear position was defined as the center of the surface-modeled nucleus (Fig. 3c, right). The outline of the visceral muscles (green) was merged with the position of the nuclei (red spheres) using image analysis software (Fig. 3c). In our previous studies relying on fixed embryos, the first indication of LR-asymmetric changes was found in nuclei in the posterior part of the AMG^20–22^. Therefore, in this study, we selected nuclei located 40-80 μm from the anterior tip of the midgut for further analysis (Fig. 3d, shown in magenta).

To detect potential defects in nuclear positioning, we measured the position of nuclei relative to the midline of the AMG. In the surface model, the midline (red) was placed along the merged points of the left and right visceral muscles at T4 (Fig. 3d). We then measured the distance from the center of each nucleus to the midline (Fig. 3e). Considering potential differences in the size of the AMG, we normalized nucleus–midline distances as a ratio (percentage) relative to the maximum width of the AMG and calculated the mean of the normalized distances in each embryo (width of the blue box in Fig. 3e). We then averaged the mean values from 10 embryos (the average number of nuclei in each embryo was 20.1±4.8) and defined this as the distance between the nuclei and the midline. Values and standard deviations were calculated for T1-T4 (Fig. 3e; 4a, b).

**Fig. 4.**
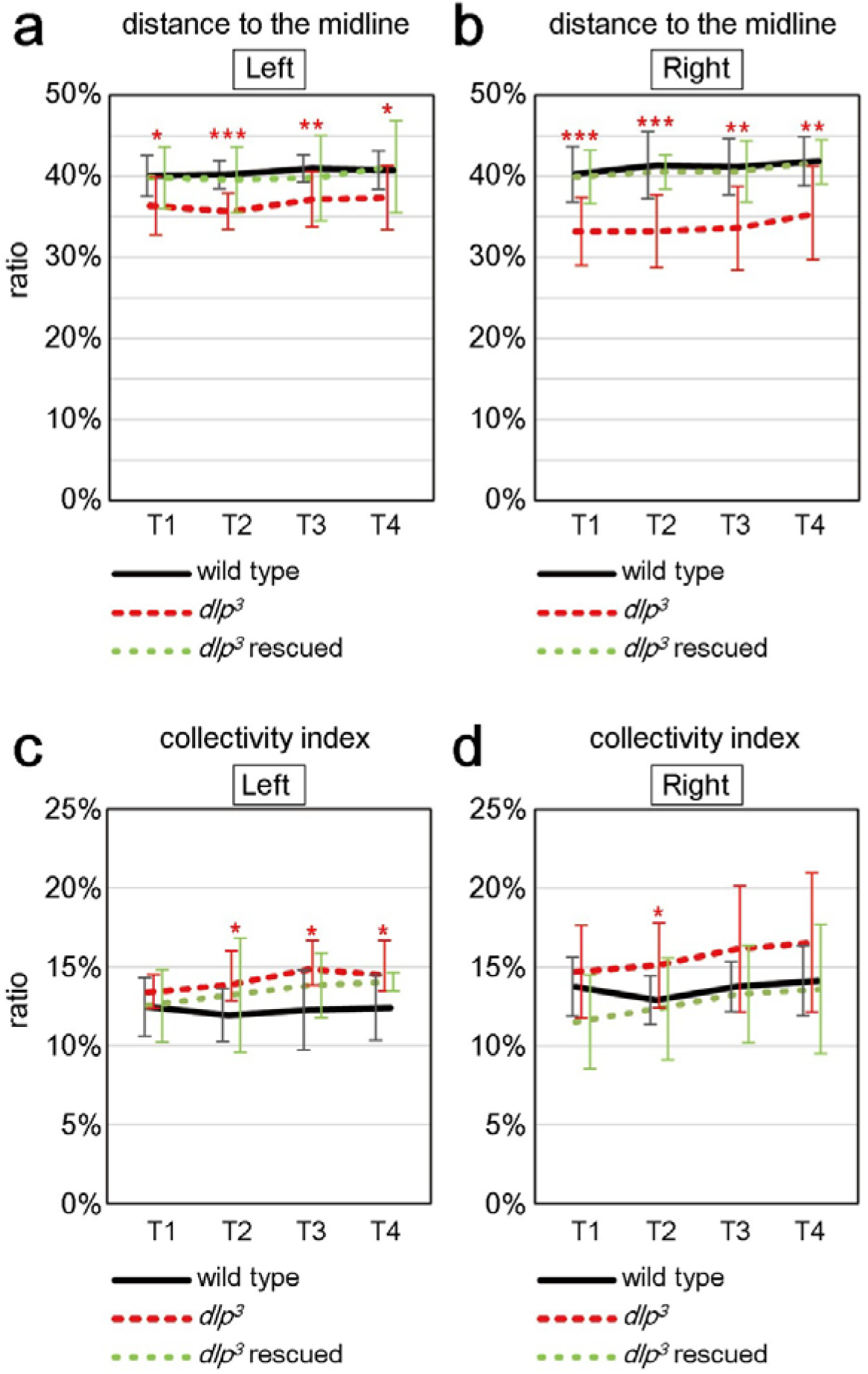
*dlp* controls the placement and collective behavior of visceral muscle nuclei. **a**, **b** The mean distance from visceral muscle nuclei to the midline, presented as a percentage of the maximum AMG width, in wild-type, *dlp^3^*, and *dlp^3^ rescued* embryos from T1 to T4. The distance to the midline, calculated as in Fig. 3e, was calculated separately for the left (**a**) and right (**b**) sides. Mean distances shown are averaged from 10 embryos. Error bars show standard deviation. **c, d** The collectivity index, which shows the mean distance between visceral muscle nuclei, was calculated as in Fig. 3f and shown as a percentage of the maximal width of the AMG, at T1 to T4. The collectivity index is calculated y averaged from the mean distance between nuclei for 10 embryos. The left (**c**) and right (**d**) sides of the AMG were analyzed separately. (**a-d**) Genotypes: wild-type (black lines), *dlp^3^* homozygotes (red), and *dlp^3^* homozygotes expressing *UAS-dlp* (green); all with *65E04-driven* expression of *UAS-RedStinger* and *UAS-lifeact-EGFP* in the visceral muscles. *p<0.05; **p<0.01; ***p<0.001.

We used this procedure to analyze the distance between the nuclei and the midline in wild-type and *dlp^3^* homozygous embryos, and found that the distance from the nucleus to the midline was significantly less in the visceral muscles of *dlp^3^* mutants, on both the right and left sides, than in wild-type embryos, at T1-T4 (Fig. 4a, b). Importantly, the specific overexpression of UAS-*dlp* in the visceral muscles, driven by *65E04*, rescued this defect in both the left and right sides in *dlp^3^* homozygotes (Fig. 4a, b). Therefore, the loss of Wnt4 signaling in the midgut visceral muscles caused mispositioning of the nuclei, such that they approached the midline more closely (on both the left and right sides) than in wild-type visceral muscles. Thus, Wnt4 signaling is required for PNP.

### Wnt4 signaling controlled the collectivity of nuclear arrangement

In 3D time-lapse movies, nuclei appeared more dispersed in the visceral muscles of *dlp^3^* mutants compared with wild-type embryos (Fig. 3a, b). To measure defects in CNB, we calculated a collectivity index to represent the mean distances between each nucleus and its nearest posterior neighbor, normalized as a percentage of the maximal width of the midgut (Fig. 3f). We then averaged the collectivity index values from 10 embryos and calculated the standard deviations at T1-T4 (Fig. 4c, d). The collectivity index of the left and right visceral muscles was higher in *dlp^3^* homozygotes than in wild-type embryos at stage T1-T4; this difference was statistically significant at T2 to T4 for the left side and at T2 for the right side (Fig. 4c, d). These results suggest that CNB depends on Wnt4 signaling. Indeed, CNB defects were rescued in the visceral muscle of *dlp*^3^ mutant embryos overexpressing *UAS-dlp*, as their collectivity index was similar to that of wild-type embryos at T2-T4 on the right side (Fig. 4d). Although the rescue effect was weaker on the left side, the collectivity index did not differ significantly between wild-type and rescued embryos (p values for T1-T4 ranged from 0.11 to 0.95) (Fig. 4c). Therefore, Wnt4 signaling in the visceral muscle regulates both PNP and CNB.

Considering our observations that the nuclei actively moved and changed their positions relative to each other in wild-type embryos (Fig. 1c, d), we speculated that the reduced collectivity of nuclei in *dlp* mutants could be due to augmented movement. Based on 3D time-lapse movies, we tracked the migration of nuclei in wild-type and *dlp*^3^ homozygous embryos by determining their position at 5-min intervals for 30 min beginning at T1 (Fig. 2f; Supplementary Figure 2f). Mean values calculated for migration distance (μm) and averaged for three embryos demonstrated that nuclei migrated farther in both left and right sides of *dlp^3^* mutants than in wild-type embryos; the difference in the right side was statistically significant (Fig. 2f). Thus, accelerated migration may be responsible for the dispersion of nuclei in *dlp* mutants.

### Myosin II and a Nesprin-like protein are required for proper positioning but not the collective behavior of the nuclei

We next examined the mechanisms underlying PNP and CNB. LINC complex, which consists of KASH- and SUN-domain proteins, physically links the nuclear envelope and the cytoskeleton and plays crucial roles in nuclear migration in several species, including *Drosophila*^24^. Muscle-specific protein 300 kDa (Msp300), a *Drosophila* KASH-domain protein (Nesprin-like protein), is required for the proper positioning of the nuclei in skeletal muscles and the eye imaginal disc^43–45^. Therefore, we investigated potential roles for *Msp300* in the LR-asymmetry of the AMG and in PNP and CNB.

We analyzed AMG LR-asymmetry using *Msp300^ΔKASH^*, an *Msp300* loss-of-function allele that encodes a mutant protein lacking the KASH domain required for its activity^44^. The predominant LR defect in *Msp3^ΔKASH^* homozygous embryos was a no-laterality phenotype in the AMG (18%) (Fig. 5a). Interestingly, surface-modeling analyses revealed that the distance between the nuclei and the midline in in the right-side visceral muscles of *Msp300^ΔKASH^* embryos at T1-T4 was significantly less in than wild-type (Fig. 5d, e). Thus, the requirement for *Msp300* in PNP was LR-asymmetric, unlike for *dlp*, which was required for both the right and left sides (Fig. 5d, e). We also analyzed CNB in *Msp300^ΔKASH^* homozygotes, and found that despite the defect in PNP, the collectivity index did not differ significantly from that of wild-type embryos at T1-T4, revealing that CNB was not markedly disrupted (Fig. 5b, c, f, g). However, the standard deviation for the right side was significantly larger in *Msp300^ΔKASH^* mutants compared to wild-type (p values: T1, 0.001; T2, 0.0005; T3, 0.01; T4. 0.03), which suggests that the collectivity index varied among the individual embryos (Fig. 5g).

**Fig. 5.**
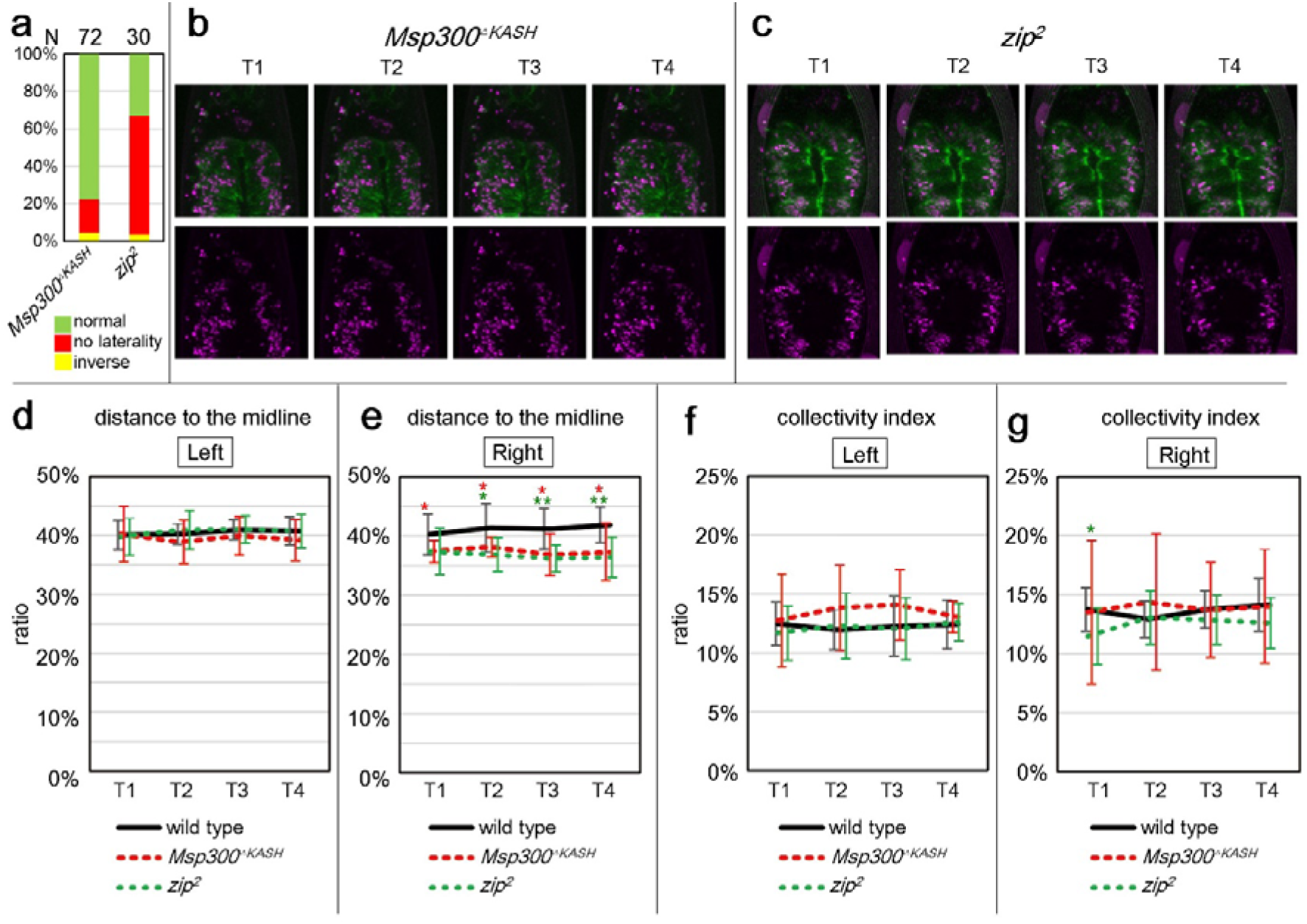
*Msp300* and *zip* are required for proper position of the nuclei but not for collective nuclear arrangement. **a** Bars show the percentage of *Msp300^ΔKASH^* and *zip^2^* homozygous embryos with LR AMG phenotypes with normal (green), no laterality (red), and inverse LR (yellow) laterality. The number of embryos examined (N) is shown above each bar. **b, c** Snapshots of 3D time-lapse images of the AMG at T1-T4, as viewed from the ventral side, in **b**) *Msp300^ΔKASH^* and **c**) *zip^2^* homozygous embryos. Top panels show nuclei (magenta) and F-actin (green); lower panels show nuclei. **d, e** The mean distance between nuclei and the midline, averaged from 10 embryos and shown as a percentage of the maximal width of the AMG, at T1 to T4. The left (**d**) and right (**e**) sides of the AMG were analyzed separately. **f, g** The collectivity index, calculated from the mean distance from nuclei to their nearest posterior neighbor and averaged from 10 embryos, are shown. The left (**f**) and right (**g**) sides of the AMG were analyzed separately. (**d-g**) Genotypes: wild-type (black), *Msp300^ΔKASH^* homozygous (red), and *zip^2^* homozygous (green) embryos, all with *65E04*-driven expression of *UAS-RedStinger* and *UAS-lifeact-EGFP* in the visceral muscles. *p<0.05; **p<0.01.

MyoII contributes to LINC complex–dependent nuclear migration in various systems by physically linking F-actin^23^. We previously reported that zipper^2^ (*zip^2^*), a mutant of the gene encoding MyoII heavy chain, produced a symmetrical AMG phenotype, reminiscent of the *Msp300^ΔKASH^* mutant phenotype, at 60% frequency (Fig. 5a)^21^. Moreover, MyoII is required in AMG visceral muscles for the organ’s normal LR-asymmetric development^21^. Given the relevance of aberrant nuclear positioning to the LR defects we observed, we analyzed CNB in *zip^2^* homozygotes. As in *Msp300^ΔKASH^* mutants, the average distance between the nuclei and the midline was decreased in *zip^2^* mutants compared to wild-type embryos at T2-T4, but only in the right-side visceral muscles (Fig. 5d, e). Thus, nuclear positioning was LR-asymmetric in *Msp300^ΔKASH^* and *zip^2^* mutants, although it was LR-symmetric in wild-type embryos (Fig. 5f, g). In other words, *Msp300* and *zip* may be required only in the right-side visceral muscles in wild-type embryos. However, the average collectivity index did not differ significantly between *zip^2^* and wild-type embryos, except for a slight reduction in the *zip^2^* mutants at T1 (Fig. 5f, g). Based on these results, we speculated that PNP is controlled by a MyoII-dependent mechanical force applied to the nuclear envelope via physical links between F-actin and the LINC complex. However, these mechanical processes may be irrelevant to CNB. Nevertheless, in mutants with defects in PNP, the LR-asymmetry of the AMG was also disturbed; but this was not always the case with defects in CNB. Therefore, PNP in the visceral muscles may be a prerequisite for establishing normal LR-asymmetry, but it may also be integral to the mechanism of LR-asymmetrical development.

## Discussion

In the developing embryo, the breaking of bilateral symmetry is the primary cue that initiates the cell signaling, gene expression, and morphological changes that support LR-asymmetric development^46–48^. In the mouse embryo, the clockwise rotation of the nodal cilia breaks bilateral symmetry by inducing the leftward flow of the extraembryonic fluid^49^. In snails and nematodes, blastomere chirality breaks the embryo’s bilateral symmetry at early cleavage stages and drives the subsequent LR-asymmetric events^1,5,11^. In these scenarios, the initial cue that initiates LR-asymmetry is gradually amplified to achieve the LR-asymmetric development of the whole body. However, our present study revealed a different strategy, in which achieving LR symmetry is a crucial step toward establishing LR-asymmetry (Fig. 6).

**Fig. 6.**
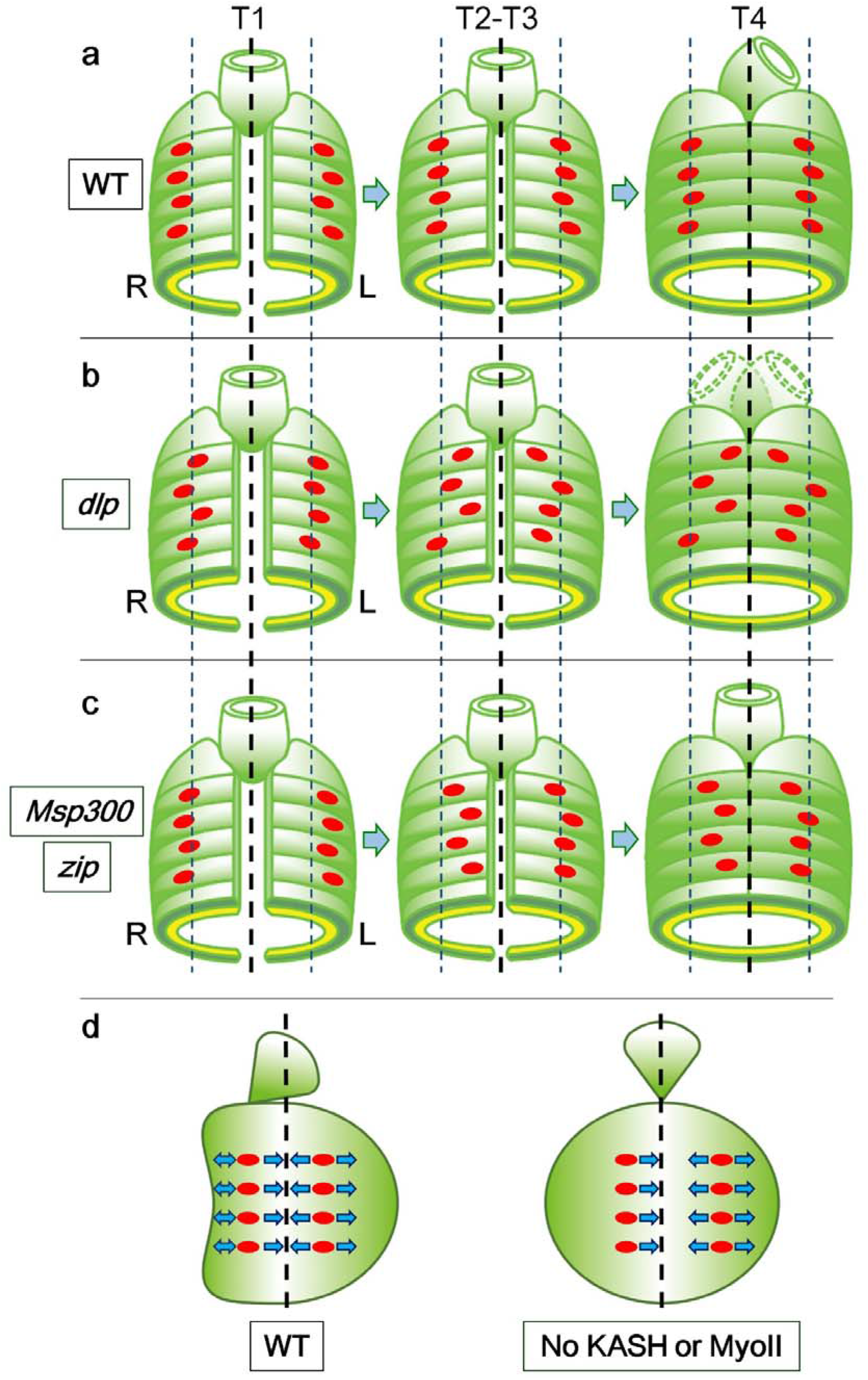
Summary of genetic controls for nuclear positioning in *Drosophila* visceral muscles. a-c Diagrams show the position of visceral muscle nuclei (red ovals) relative to the midline (dotted black line) at T1 (left), T2-T3 (middle), and T4 (right). Dotted blue lines show normal nuclear position. R, right; L, left. **a** In wild-type embryos, the nuclei align in a rib-cageshaped zone along the anterior–posterior axis in both the right and left: sides of the AMG, and can move laterally relative to each other. The nuclei become LR-symmetrically aligned by T4, when the AMG starts its LR-asymmetric development. **b** Nuclei in *dlp^3^* homozygotes are more dispersed and are closer to the midline than those in wild-type embryos. Under these conditions, the LR-asymmetry of the AMG becomes randomized. **c** In *Msp-300* or *zip* mutants, the right-side visceral muscle nuclei are positioned closer to the midline than in wild-type embryos. Thus, LINC complex and MyoII play an LR-asymmetric role in preserving the distance between the nuclei and the midline. In contrast to *dlp* mutants, nuclei n *Msp-300* and *zip* mutants retain the ability to behave collectively. In these mutants, the AMG remains LR-symmetric at T4 and afterwards. **d** A model demonstrating the role of bilaterally asymmetric nuclear positioning in the LR-asymmetric morphogenesis of the AMG. The LR-symmetric pulling force acting on nuclei on the right side may be coupled with LR-asymmetric morphogenesis.

Here, we demonstrated that the bilaterally symmetric arrangement of the nuclei in the visceral muscles of the AMG is required for this organ’s LR-asymmetric development (Fig. 6). In the absence of MyoII or a LINC-complex component, the nuclei align LR-asymmetrically but the AMG develops LR-symmetrically (Fig. 6a, c). Thus, MyoII and LINC complex play important roles in the LR-symmetric rearrangement of the nuclei, which is required for or coupled with the subsequent LR-asymmetric morphogenesis (Fig. 6d). We speculate that such translation between lateral symmetry and asymmetry may act as an additional layer of regulatory steps, and that this multi-layered regulation allows multiple mechanisms to contribute to LR-asymmetric development in a species. In such cases, complex LR-asymmetric structures can be built with a limited number of machineries.

Although a requirement for proper nuclear positioning in LR-asymmetry has not been reported previously, defective nuclear positioning has been connected to human diseases^27–28^. Mutations of genes that encode key molecules for nuclear positioning, such as LINC complex, are associated with Emery-Dreifuss muscular dystrophy and cerebellar ataxia^23^, and genetic analysis in model animals revealed that LINC complex plays key roles in the development of these diseases^28^. In *Drosophila* optic epithelium and in vertebrate neuroectoderm, defects in nuclear migration and positioning affect the pattern of cell division^43,50^. However, it is unlikely that cell division initiates the LR-asymmetric development of the AMG, because cell propagation is complete before the collective nuclear rearrangement and LR-asymmetric development of this organ begins^51^. On the other hand, defective nuclear positioning may mechanically influence AMG morphogenesis. The nucleus can act as a piston that physically compartmentalizes the cytoplasm and provides hydrostatic pressure toward the direction of nuclear migration^24,52^. Given that the nucleus can provide this type of dynamic force, the positioning of the nuclei may help to create mechanical forces that promote LR-asymmetric morphogenesis.

Here we revealed two distinct events that control nuclear location PNP and CNB (Fig. 6a, b, c), both of which require Wnt4 signaling (Fig. 6b). However, LINC complex and MyoII are required for PNP but not for CNB (Fig. 6c), demonstrating that the two events depend on distinct underlying mechanisms. MyoII provides contractile force to F-actin, which is involved in LINC complex–dependent nuclear movement in other systems^23^. Considering that nuclei in the right-side visceral muscles shifted toward the midline in the absence of MyoII or a LINC complex component, these two factors may introduce an ability to resist a pulling force from the midline (Fig. 6d). Resistance to such a pulling force might also derive from the counteracting forces of LR-asymmetric tissue deformation (Fig. 6d, e). This idea is consistent with our observation that AMG morphogenesis was bilaterally symmetrical in the absence of MyoII or a LINC complex component.

We also found that Wnt4 signaling is required for the CNB in the visceral muscles (Fig. 6b). In the wild-type embryo, the nuclei are densely packed into a limited area in each lateral half of the ventral region of the AMG (Fig. 6a). However, when Wnt4 signaling was interrupted, as in *dlp* mutants, the nuclei were sparsely distributed over a larger area and migrated more actively (Fig. 2f and Fig. 6b). This observation suggests that Wnt4 signaling might organize the collective movement of the nuclei in wild-type embryos by downregulating nuclear migration. A specific association between the LR-randomization phenotype and defects in Wnt4 signaling suggests that defective nuclear placement, shown by a more dispersed distribution and a failure to preserve a distance from the midline, might contribute to LR randomization (Fig. 6b)^22^. If this is the case, proper placement of the nuclei may be important for directing the LR polarization of the mechanical force driving AMG morphogenesis. Our results also suggest that the degree of the CNB varied between individual *Msp300^ΔKASH^* mutants, even though their collectivity index did not differ significantly from that of wild-type embryos. However, *Drosophila* has multiple KASH genes, and their redundant functions may make it difficult to fully ascertain *Msp300’s* roles in CNB^53^.

In this analysis, we demonstrated that nuclear position is crucial in forming LR-asymmetry. Considering that non-skeletal muscles—which are, like *Drosophila* visceral muscles, formed of multi-nucleated cells—contribute to LR-asymmetric organs and tissues such as the heart, blood vessels, and digestive organs in vertebrates and other organisms, the contribution of nuclear positioning to LR-asymmetric development may be evolutionarily conserved.

## Methods

### *Drosophila* stock

We used Canton-S as the wild-type (WT) control strain. We also used *Drosophila* lines with the following genotypes: *dlp^3^*, a loss-of-function allele (induced by ethyl methane sulfonate in this study); *dlp^MH20^*, a null allele^54^; *zip^2^*, an amorphic allele^55^; and *Msp300^ΔKASH^*, a loss-of-function allele^44^. We used the UAS lines *UAS-lifeact-EGFP*^32^, *UAS-Redstinger*^56^, *UAS-dlp*^57^, and *UAS-dshp*^22^, and the following Gal4-driver lines: *arm-Gal4*, which is an ubiquitous driver^58^; *hand-Gal4*^59^ and *65E04*^60^, which are specific to the circular visceral muscle; *24B*, which is specific to the somatic, circular visceral, and longitudinal visceral muscles^61^; *NP5021*, which is specific to the endodermal epitheliumr^62^; and *Elav-Gal4*, a pan-neuronal driver^63^.

Mutations on the second chromosome were balanced with *Cyoβ*. Mutations on the third chromosome were balanced with *TM2β*. All genetic crosses were performed at 25°C on standard *Drosophila* culture media.

### Immunostaining

Immunostaining of embryos was as previously described^64^. We used the following primary antibodies: mouse anti-β-galactosidase (Promega, 1:1,000 dilution), rabbit anti-GFP (MBL,1:1,000 dilution), rat anti-HA (3F10; Roche Diagnostics, 1:1,000 dilution), mouse anti-Connectin [C1.427, Developmental Studies Hybridoma Bank (DSHB), 1:5 dilution], mouse anti-Crumbs (Cq4, DSHB, 1:30 dilution), and mouse anti-FasIII (7G10, DSHB, 1:100 dilution). We used Cy3-conjugated anti-mouse IgG (Jackson ImmunoResearch, 1:500) and biotinylated anti-mouse IgG (Vector Labs, 1:200 dilution) as secondary antibodies. We used the Vectastain ABC kit (Vector Labs) for biotin-staining reactions.

### Analysis of LR-asymmetry in the AMG

Embryos were fixed and stained with anti-Fas3 (DSHB, 1:50) as described previously^22^. Images were obtained with an LSM 700 scanning laser confocal microscope (Carl Zeiss). The LR-asymmetry of the AMG in the fixed embryo was scored based on the position of the joint between the proventriculus and the AMG relative to the midline, as previously described^22^. Briefly, if the joint was to the left of the midline, the phenotype was scored as normal; if to the right, it was scored as inverse, and if overlapping the midline, as no laterality.

### Live imaging

Embryos with the following genotypes were collected before stage 13: *65E04, UAS-redstinger/UAS-lifeact-EGFP* (used as wild-type); 65E04-GAL4,dlp3,UAS-redstinger/65E04-GAL4 dlp3,UAS-lifeact-EGFP-p10 (used as *dlp^3^* mutant); *UAS-RedStinger/+;dlp^MH20^, 65E04-gal4,UAS-Lifeact-GFP/UAS-dlp, dlp^MH20^* (used as *dlp^3^* rescued); *Msp300^ΔKASH^/Msp300^ΔKASH^*; 65E04, UAS-redstinger/65E04, UAS-lifeact-EGFP (used as *Msp300^ΔKASH^* mutant); and *zip^2^/zip^2^; 65E04*, UAS-redstinger/65E04, UAS-lifeact-EGFP (used as *zip^2^* mutant). Embryonic eggshells were removed by immersion in 50% bleach for 1 min, followed by a wash in water. Stage 13 embryos of the appropriate genotype were selected under florescence microscopy, mounted ventral-side up on double sticky tape on a glass slide, placed between 0.25 mm-thick spacers made from coverslips, mounted in oxygen-permeable Halocarbon oil 27 (Sigma), and covered with a coverslip^3^. 3D time-lapse movies of embryos at 18°C were taken every 10 min for 2 hr using the LSM 880 (Carl Zeiss). At stage 13, the anterior–posterior axis of the embryo (identified by head and tail structures), was manually reoriented to the Y-axis of the image. We obtained 3D time-lapse movies using Z stacks (13-15 images at 5-μm intervals). To reduce phototoxicity, we used a relatively low laser power (488 nm laser, 0.03-1; 561 nm laser, 0.3-5). The time-lapse images were saved as LSM files in ZEN software (2012 SP1 black edition, Release Version 8.1, Carl Zeiss).

### 3D reconstruction of nuclear movement in the visceral muscles of the AMG

The time-lapse movies were 3D-reconstructed using Imaris image analysis software (Bitplane). The positions of the surface-modeled nuclei in 3D coordinates were determined for each time point using the Spot function. The 3D surface models of the visceral muscles were constructed using the Surface function. The 3D surface models and the positions of the center of each nucleus were saved as VRML (Virtual Reality Modeling Language) files, to be analyzed simultaneously.

### File translation to construct the 3D-surface model

To easily obtain measurements in 3D space, Maya version 2018 (Autodesk, San Rafael, CA), a computer animation and modeling software, was used to animate the 3D-surface model^65^. To use Maya, the VRML files, in which the 3D-surface models of visceral muscles and the centers of the nuclei were integrated, were converted to 3DS (native file format of the old Autodesk 3D Studio DOS) using Meshconv (https://www.patrickmin.com/meshconv).

### Preprocessing the surface models

The 3DS files were imported into Maya version 2018 (Autodesk). The surface models of the visceral muscles were transparently colored and then added to the layers (Fig. 3c). Although *65E04*-driven *UAS-Redstinger* expression is highly specific to the visceral muscles, some nuclei outside the visceral muscles were also labeled. Therefore, any nuclei that lay outside the surface-modeled visceral muscles were manually deleted. In previous studies, we detected the LR-asymmetric rearrangement of the nuclei in the posterior half of the AMG at stage 15.

Thus, in this study, the nuclei were divided by whether they were in the anterior or posterior region of the AMG, corresponding to 0-40 μm and 40-80 μm from the anterior tip of the midgut, respectively (Fig. 3d), and nuclei in the posterior region were analyzed further.

### Measuring the average distance between the nuclei and the midline

Distances between the nuclei and the midline of the AMG were measured using Maya version 2018 (Autodesk)^65^. To define the midline, we used Maya’s Convert function to fit the surface-modeled AMG in a minimal cuboid placed along the anterior–posterior axis of each embryo (Fig. 3d). Thus, the width of the cuboid corresponded to the maximal width of the AMG. The midline was manually placed in the 3D-surface model as a line parallel to the anterior–posterior grids of the cuboid that connected the points where the left- and right-side visceral muscles merge at T4 (Fig. 3d). This midline was used retrospectively for data captured at T1 to T3.

We measured distances between nuclei in the posterior part of the surface model (40-80 μm from the anterior tip) and the midline by measuring the length of the line drawn perpendicularly from the midline to the nucleus (Fig. 3e). To adjust for differences in the size of the AMG, the values were normalized as a ratio (percentage) of the maximum width of the cuboid, and the mean of the normalized distances was calculated for each embryo. This procedure was done automatically using Python Script in Maya and NumPy library (Supplement Figure, script). We then averaged the mean values and calculated standard deviations for T1-T4.

### Measuring the collectivity of nuclear arrangement

To analyze the collectivity of nuclear arrangement, we measured the distance between each nucleus and its nearest-most-posterior neighbor, and calculated the mean distance for each embryo. We used Python Script and NumPy library in Maya version 2018 (Autodesk), to automatically connect each nucleus in the lower region of the surface model (40-80 μm below the anterior tip) to its nearest most-posterior neighbor and to calculate the length of the connecting line (Fig. 3f). Distances were measured separately for the right and left sides of the visceral muscles, and mean values were obtained for each embryo (Fig. 3f). We averaged the mean values and calculated standard deviations for T1-T4. To normalize differences in embryo size, values are presented as a percentage of the maximum width of the AMG (the width of the cuboid) (Fig. 3f).

### Measuring the migration path of the nuclei

We measured migration distance by tracing the paths traveled by the nuclei. We used 3D time-lapse movies obtained at 5-min intervals over a 30-min period. The 3D movies, composed of 13-15 Z stacks, were converted to 2D-sequence image files using the maximum intensity projection feature in ZEN software (Carl Zeiss). 2D-sequence image files were imported into Maya version 2018 (Autodesk) and displayed to track the migrating nuclei. We manually traced the path of each nucleus through the sequence of images using Maya’s EP curve tool. Then, the length of each traced path (μm) was automatically measured using a Python script in Maya (See Supplement). From these measurements, we calculated the average migration distance and standard deviation.

### Statistical processing

Statistical processing was done in Maya version 2018 (Autodesk) and Excel 2013 (Microsoft, Redmond, WA). The calculated values were copied to Excel, and the average of the mean values and their standard deviations were calculated using the AVERAGE and STDEVP functions. To evaluate the statistical significance of the differences between phenotypes, we used Excel’s F.TEST and T.TEST functions.

## Supporting information

Raw data for Fig 5. Msp300 and zip are required for proper nuclear placement but not for collective nuclear arrangement.

Raw data for Fig 4. dlp controls nuclear placement and collectivity.

Graph and scripts

## Notes

### Competing Interest Statement

The authors have declared no competing interest.

