## Supplementary material for "Collective nuclear behavior shapes bilateral nuclear symmetry for subsequent left-right asymmetric morphogenesis in *Drosophila*": Graph and scripts

### Slide 1
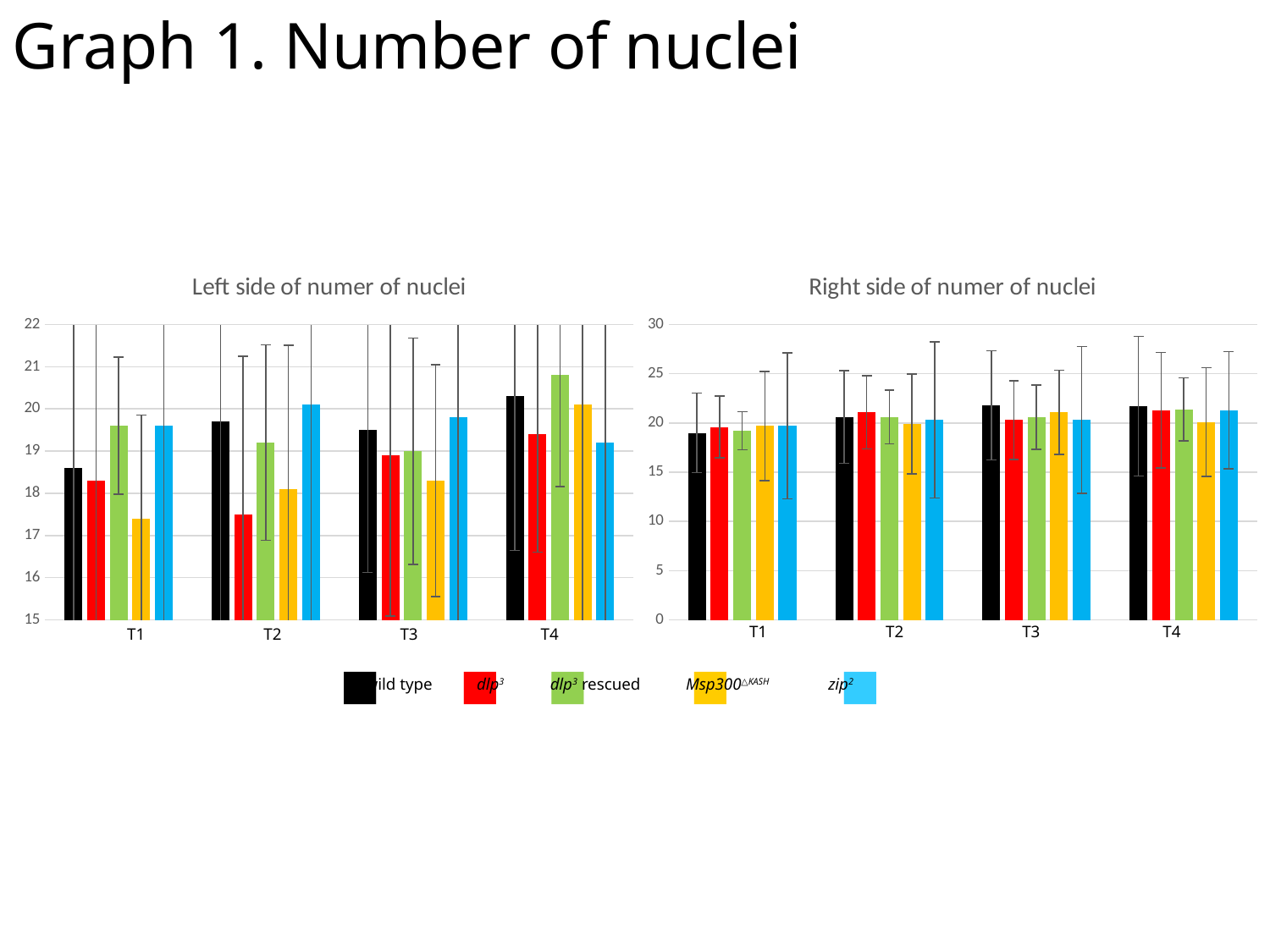

Graph 1. Number of nuclei
###### Chart: Left side of numer of nuclei
| Category | wt_av | dlp_av | rescue_av | kash_av | zip_av |
|---|---|---|---|---|---|
###### Chart: Right side of numer of nuclei
| Category | wt_av | dlp_av | rescue_av | kash_av | zip_av |
|---|---|---|---|---|---|T1 T2 T3 T4
T1 T2 T3 T4
wild type
dlp^3^
dlp^3^ rescued
Msp300^△KASH^
zip^2^
■ ■ ■ ■ ■

### Slide 2
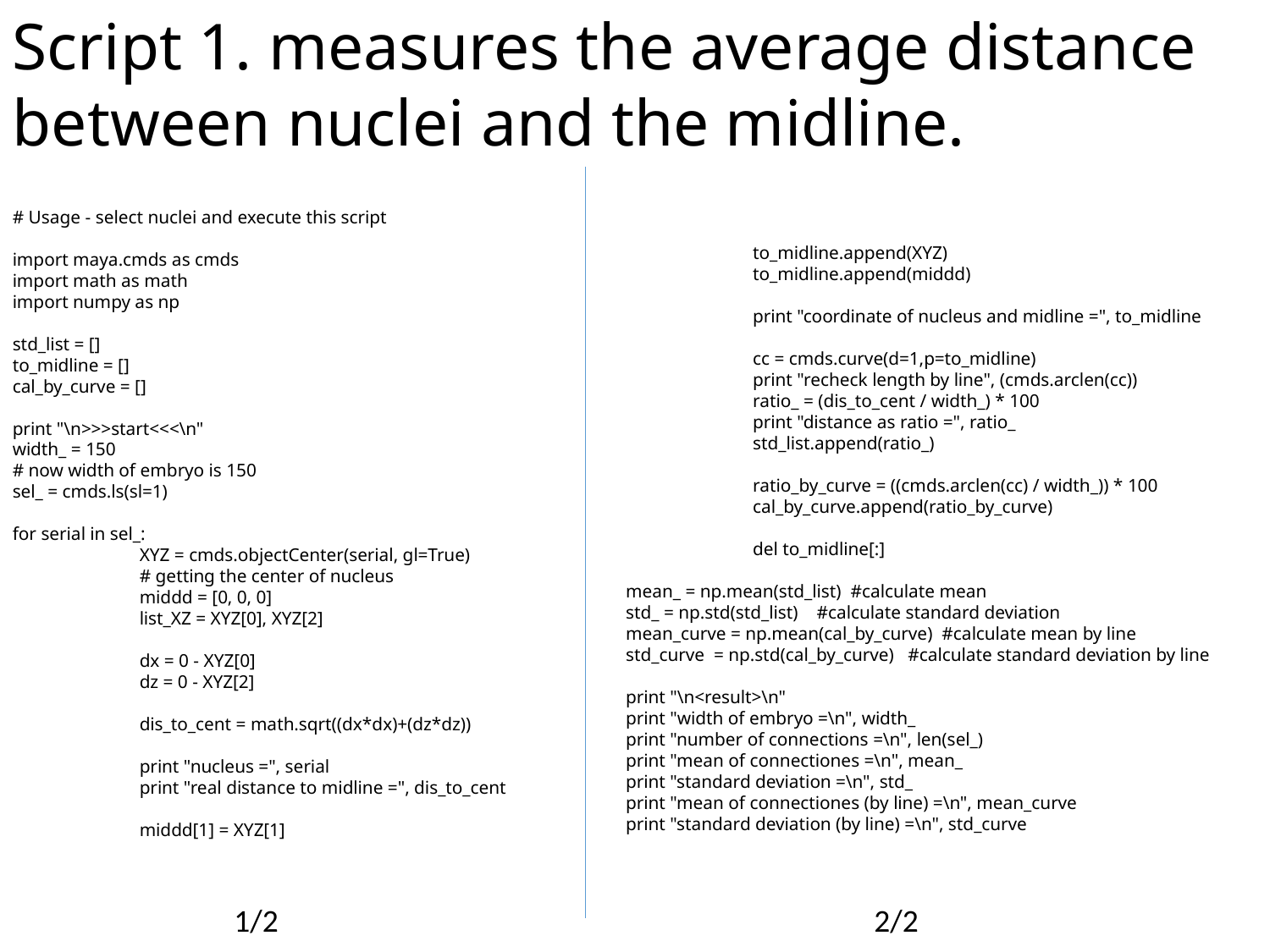

Script 1. measures the average distance between nuclei and the midline.
#### Usage - select nuclei and execute this script
import maya.cmds as cmds
import math as math
import numpy as np
std_list = []
to_midline = []
cal_by_curve = []
print "\n>>>start<<<\n"
width_ = 150
#### now width of embryo is 150
sel_ = cmds.ls(sl=1)
for serial in sel_:
	XYZ = cmds.objectCenter(serial, gl=True)
	# getting the center of nucleus
	middd = [0, 0, 0]
	list_XZ = XYZ[0], XYZ[2]
	dx = 0 - XYZ[0]
	dz = 0 - XYZ[2]
	dis_to_cent = math.sqrt((dx*dx)+(dz*dz))
	print "nucleus =", serial
	print "real distance to midline =", dis_to_cent
	middd[1] = XYZ[1]
	to_midline.append(XYZ)
	to_midline.append(middd)
	print "coordinate of nucleus and midline =", to_midline
	cc = cmds.curve(d=1,p=to_midline)
	print "recheck length by line", (cmds.arclen(cc))
	ratio_ = (dis_to_cent / width_) * 100
	print "distance as ratio =", ratio_
	std_list.append(ratio_)
	ratio_by_curve = ((cmds.arclen(cc) / width_)) * 100
	cal_by_curve.append(ratio_by_curve)
	del to_midline[:]
mean_ = np.mean(std_list) #calculate mean
std_ = np.std(std_list) #calculate standard deviation
mean_curve = np.mean(cal_by_curve) #calculate mean by line
std_curve = np.std(cal_by_curve) #calculate standard deviation by line
print "\n<result>\n"
print "width of embryo =\n", width_
print "number of connections =\n", len(sel_)
print "mean of connectiones =\n", mean_
print "standard deviation =\n", std_
print "mean of connectiones (by line) =\n", mean_curve
print "standard deviation (by line) =\n", std_curve
1/2 2/2

### Slide 3
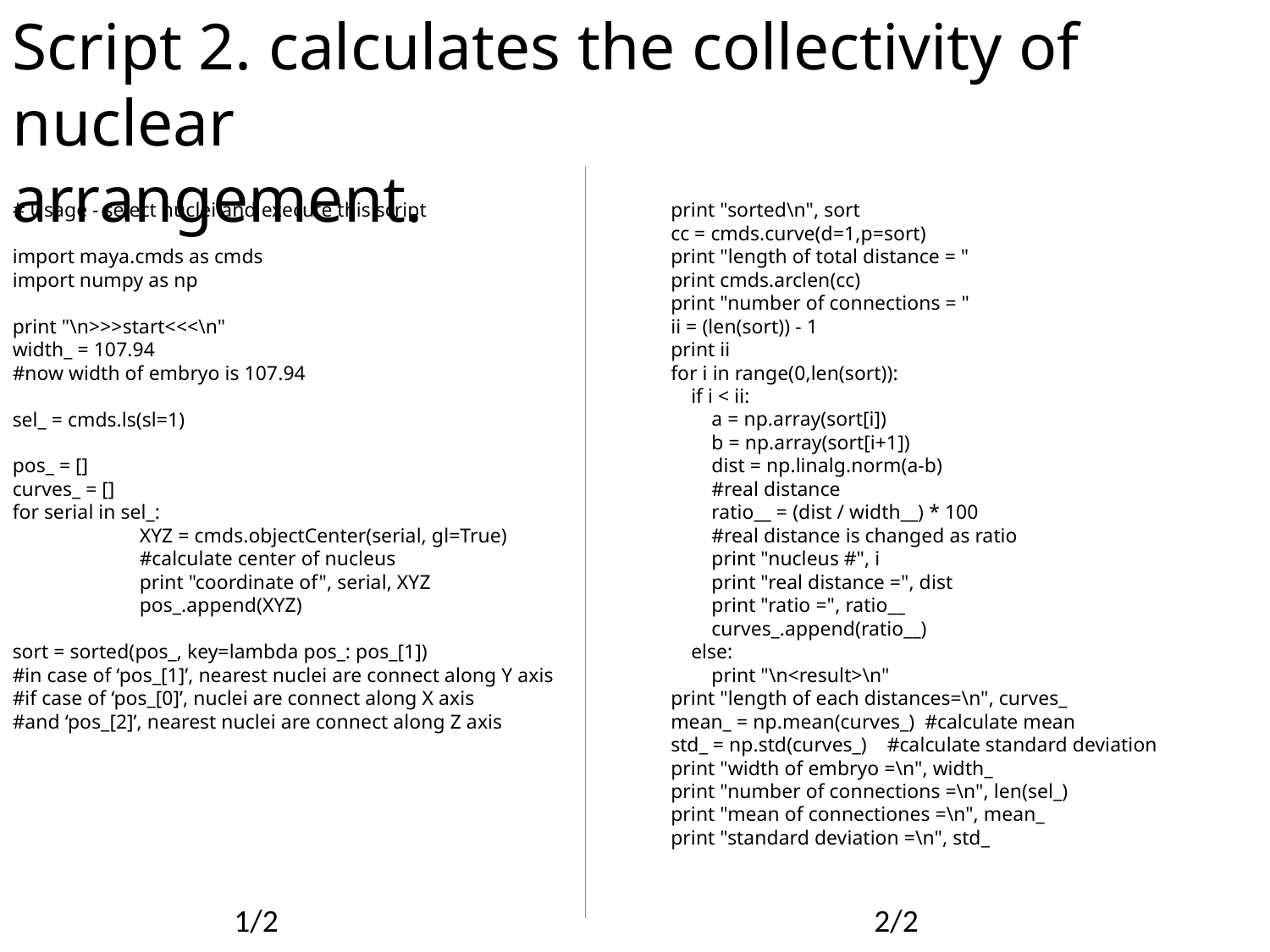

Script 2. calculates the collectivity of nuclear
arrangement.
#### Usage - select nuclei and execute this script
import maya.cmds as cmds
import numpy as np
print "\n>>>start<<<\n"
width_ = 107.94
#now width of embryo is 107.94
sel_ = cmds.ls(sl=1)
pos_ = []
curves_ = []
for serial in sel_:
	XYZ = cmds.objectCenter(serial, gl=True)
	#calculate center of nucleus
	print "coordinate of", serial, XYZ
	pos_.append(XYZ)
sort = sorted(pos_, key=lambda pos_: pos_[1])
#in case of ‘pos_[1]’, nearest nuclei are connect along Y axis
#if case of ‘pos_[0]’, nuclei are connect along X axis
#and ‘pos_[2]’, nearest nuclei are connect along Z axis
print "sorted\n", sort
cc = cmds.curve(d=1,p=sort)
print "length of total distance = "
print cmds.arclen(cc)
print "number of connections = "
ii = (len(sort)) - 1
print ii
for i in range(0,len(sort)):
 if i < ii:
 a = np.array(sort[i])
 b = np.array(sort[i+1])
 dist = np.linalg.norm(a-b)
 #real distance
 ratio__ = (dist / width__) * 100
 #real distance is changed as ratio
 print "nucleus #", i
 print "real distance =", dist
 print "ratio =", ratio__
 curves_.append(ratio__)
 else:
 print "\n<result>\n"
print "length of each distances=\n", curves_
mean_ = np.mean(curves_) #calculate mean
std_ = np.std(curves_) #calculate standard deviation
print "width of embryo =\n", width_
print "number of connections =\n", len(sel_)
print "mean of connectiones =\n", mean_
print "standard deviation =\n", std_
1/2 2/2
